# NicheNet-Based Dissection of CD14^+^ Monocyte Crosstalk with Memory CD4^+^ T Cells in Human PBMCs

**DOI:** 10.1101/2025.08.26.672385

**Authors:** Tharun Kota

**Affiliations:** university college dublin

**Keywords:** Peripheral blood mononuclear cells (PBMCs), CD14^+^ monocytes, Memory CD4^+^ T cells, Cell–cell communication, Ligand–receptor interactions, NicheNet, Immune regulation, Cytokine signaling

## Abstract

Intercellular communication between monocytes and T cells is essential for immune regulation within peripheral blood mononuclear cells (PBMCs). To examine how CD14^+^ monocytes regulate memory CD4^+^ T cells at the gene expression level, we applied NicheNet to PBMC single-cell RNA-seq data. We performed quality control, normalization, principal component analysis, and UMAP-based visualization. CD14^+^ monocytes and memory CD4^+^ T cells were identified by canonical markers and designated as sender and receiver populations, respectively. NicheNet analysis revealed both classical cytokine signaling and checkpoint-like interactions. Prominent ligands included TNF, IFNG, IL10, and CXCL9, predicted to regulate inflammatory target genes through receptors such as TNFRSF1A/B and CXCR3. In addition, the SIRPG–CD47 axis emerged as a potential costimulatory pathway modulating T cell responses. These results suggest that monocytes regulate memory CD4^+^ T cells via a combination of inflammatory, costimulatory, and regulatory signals, providing mechanistic hypotheses for immune modulation in health and disease.

## 1. Introduction

Intercellular communication underpins the coordination of immune responses, particularly within peripheral blood mononuclear cells (PBMCs), a heterogeneous mixture of T cells, B cells, monocytes, natural killer (NK) cells, and dendritic cells (Guilliams et al., 2018). These cells interact dynamically through ligand–receptor signaling, cytokine secretion, and antigen presentation, shaping immune function under both homeostatic and inflammatory conditions. Among these, CD14^+^ monocytes serve as professional antigen-presenting cells that bridge innate and adaptive immunity by processing antigens and producing cytokines and chemokines (Mosser and Edwards, 2008). Through these mechanisms, monocytes provide signals that influence the activation, differentiation, and survival of T lymphocytes (O’Shea and Murray, 2008).

Memory CD4^+^ T cells play a particularly important role in adaptive immunity, as they retain antigen specificity from prior encounters and can mount rapid and robust responses upon re-exposure to pathogens (Kumar et al., 2018). These cells are especially sensitive to cues from monocytes and other antigen-presenting cells, which help fine-tune their effector functions. Chemokine pathways also regulate this interaction, with CXCR3 and its ligands orchestrating T cell trafficking and effector responses during immune challenges (Groom and Luster, 2011). Despite extensive knowledge of cytokine and chemokine biology, the precise ligand–receptor interactions driving monocyte-mediated regulation of memory CD4^+^ T cells remain incompletely understood, particularly at the transcriptomic level.

To address this gap, we employed NicheNet, a computational framework that integrates single-cell RNA sequencing (scRNA-seq) data with curated ligand–receptor and gene regulatory networks to infer intercellular communication pathways (Browaeys et al., 2020). By focusing on PBMCs, we aimed to systematically identify which ligands expressed by CD14^+^ monocytes are most likely to drive transcriptional changes in memory CD4^+^ T cells. This approach enabled us to reconstruct ligand–receptor–target gene pathways and generate mechanistic hypotheses about how monocytes influence T cell behavior, thereby providing insights into immune regulation in human peripheral blood.

## 2. Methodology

### 2.1. Data source

We utilized the publicly accessible PBMC single-cell RNA sequencing dataset available within the Seurat R package, which includes multiple canonical immune subsets such as monocytes and T cells (Hao et al., 2021).

### 2.2. Preprocessing and Quality Control

Data preprocessing was carried out in Seurat v4.0 using standard quality control filters: only cells with 200–2,500 detected features and less than 5% mitochondrial gene content were retained. Normalization was performed with the LogNormalize method (scale factor = 10,000), followed by identification of highly variable genes.

### 2.3. Dimensionality Reduction

Dimensionality reduction was carried out using PCA on variable features. The elbow plot method (ElbowPlot in Seurat) guided the selection of the number of PCs by plotting the explained variance per component and identifying the point of diminishing returns. According to this heuristic, we selected the first 10 PCs that sufficiently summarized the dataset while minimizing overfitting (Butler et al., 2018). These PCs were then used for UMAP-based visualization of cellular heterogeneity.

### 2.4. Cell Type Annotation

Immune cell subsets were identified on the basis of canonical marker gene expression. CD14^+^ monocytes were defined by high expression of CD14 and LYZ, consistent with their established role as classical monocyte markers (Villani et al., 2017). Memory CD4^+^ T cells were characterized by expression of IL7R, CCR7, and S100A4, markers widely associated with antigen-experienced T cell populations (Zhang et al., 2005);(Mahnke et al., 2013). For the purposes of NicheNet analysis, CD14^+^ monocytes were designated as the sender population, reflecting their capacity to secrete ligands, whereas memory CD4^+^ T cells were defined as the receiver population, responding to these external cues through receptor-mediated signaling.

### 2.5. NicheNet Analysis Workflow

The NicheNet pipeline was carried out using the nichenetr R package to characterize ligand–receptor–target interactions between CD14^+^ monocytes and memory CD4^+^ T cells.

First, a target gene set was defined by performing differential expression analysis within the receiver population of memory CD4^+^ T cells, yielding candidate genes of interest. Next, candidate ligands were identified from genes expressed in CD14^+^ monocytes and cross-referenced with NicheNet’s curated ligand database. Expressed receptors on memory CD4^+^ T cells were then matched with these ligands to establish potential points of communication. To prioritize signals, ligand activity scores were calculated based on each ligand’s ability to predict the observed transcriptional program in the receiver cells (Browaeys et al., 2020). Subsequently, weighted ligand–receptor and ligand–target interaction networks were generated using NicheNet functions, applying a correlation cutoff of 0.33 to filter for robust interactions. Finally, the predicted communication pathways were summarized and interpreted through visualizations such as chord diagrams and heatmaps, providing a systems-level view of monocyte–T cell signaling.

## 3. Results

### 3.1. Ligand Activity Distribution

To evaluate the overall activity of candidate ligands expressed by CD14^+^ monocytes, we first assessed their Pearson correlation coefficient (PCC) scores with the target gene set in memory CD4^+^ T cells. As shown in Figure 1, the distribution of ligand activity scores was centered close to zero, with only a small subset of ligands surpassing the significance threshold (red dashed line). This indicates that while many ligands were expressed, only a fraction demonstrated meaningful predictive power for receiver cell gene expression changes.

**Figure 1:**
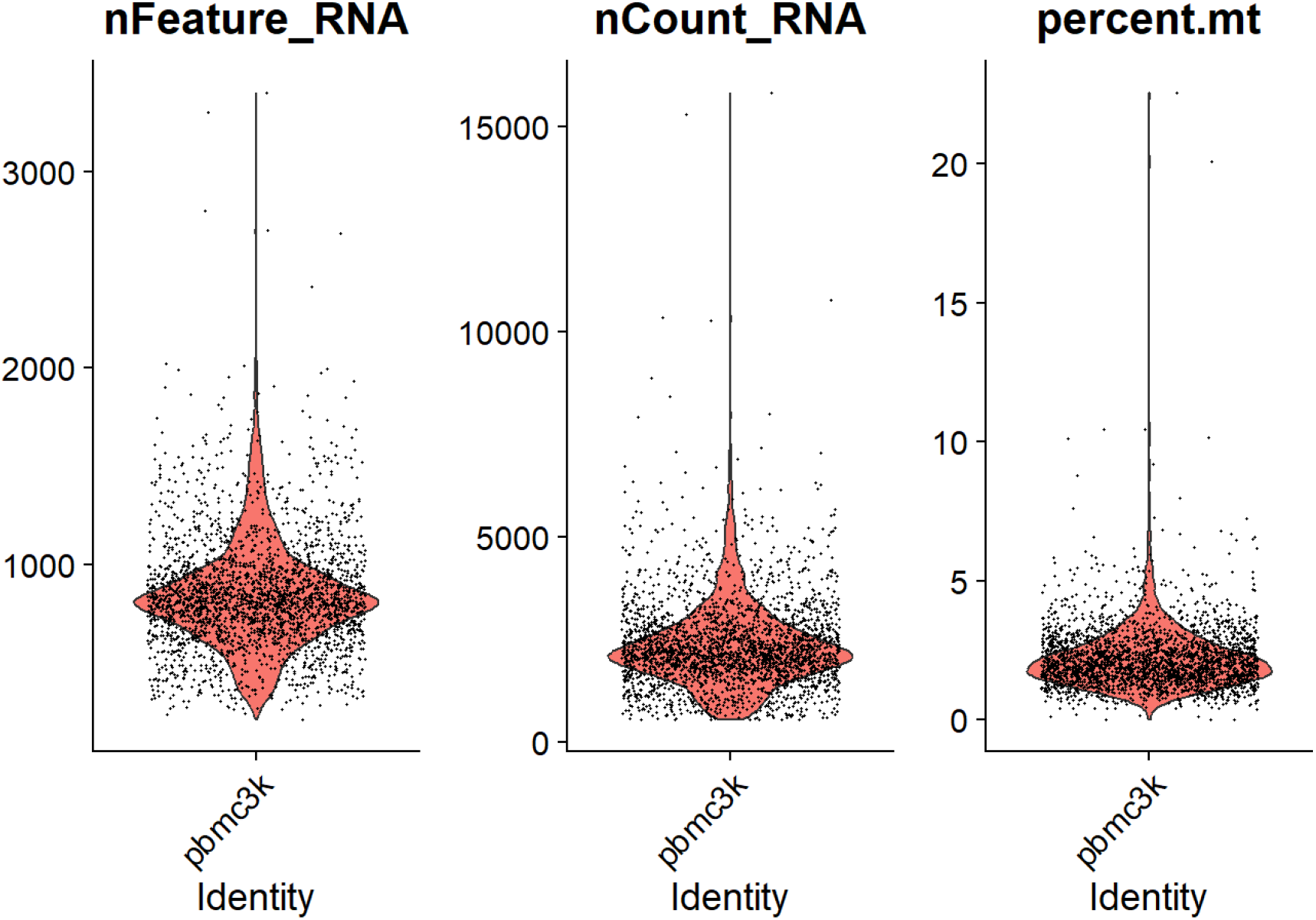
Quality control metrics of PBMC3k dataset. Violin plots showing the distribution of (A) number of detected genes per cell (nFeature RNA), (B) total RNA molecules per cell (nCount RNA), and (C) percentage of mitochondrial gene counts (percent.mt). These QC metrics are used to identify low-quality or stressed cells, with thresholds typically set to retain cells with 200–2,500 genes detected and ¡5% mitochondrial content.

### 3.2. Prioritized Ligands

NicheNet ranked ligands according to their activity scores (area under the precision-recall curve, AUPR). The top ligands included SIRPG, UCN2, SIRPB2, CD47, and CXCL9, along with classical immune mediators such as CD40LG, TNF, IFNG, and IL10 (Figure 2). These ligands represent potential drivers of gene regulation in memory CD4^+^ T cells, suggesting that both co-stimulatory signals (e.g., CD40LG) and inflammatory cytokines (e.g., TNF, IFNG) contribute to intercellular communication in this system.

**Figure 2:**
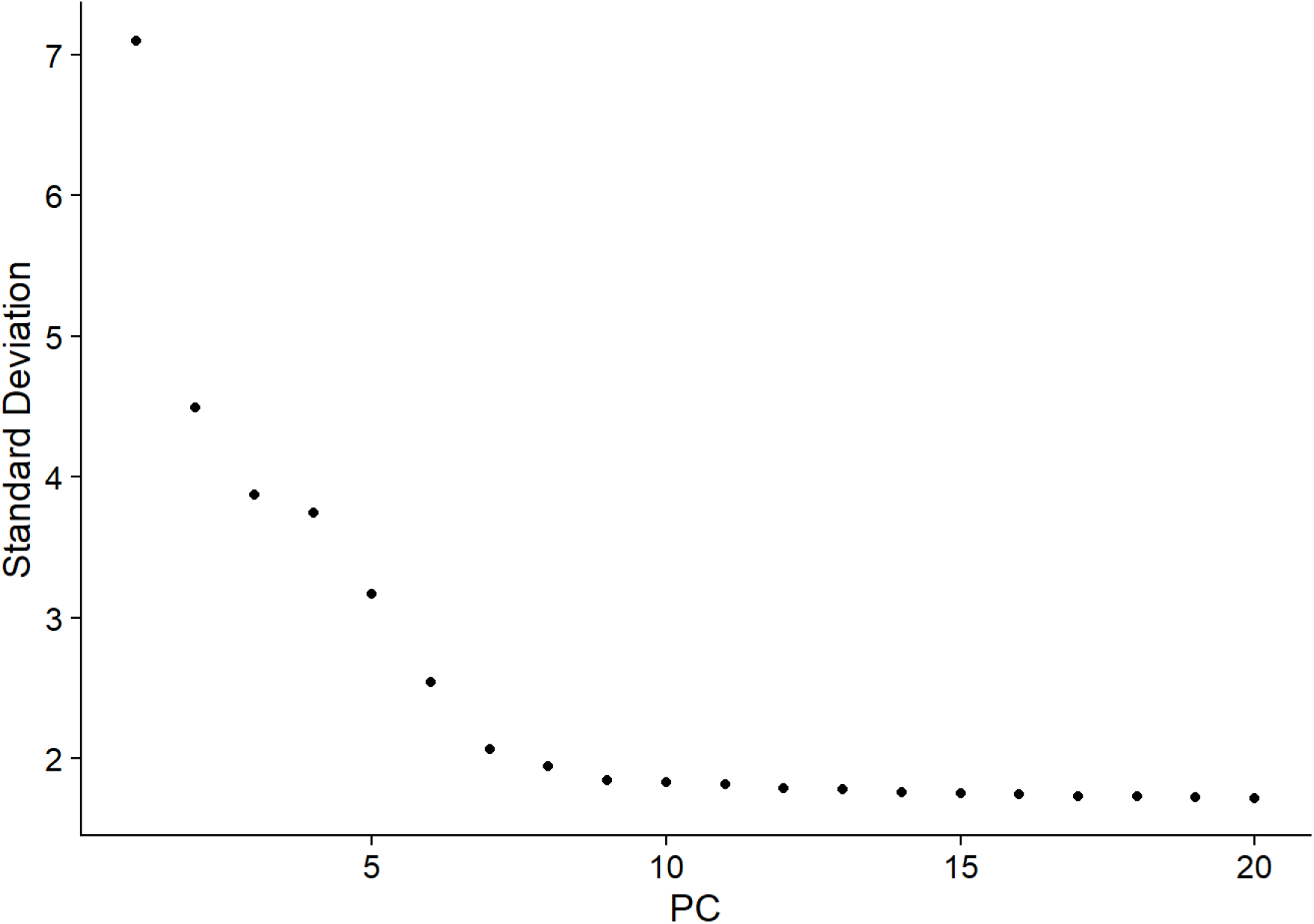
Elbow plot for principal component selection. Standard deviations of the first 20 principal components (PCs) are shown. The inflection point (“elbow”) indicates where additional PCs contribute diminishing variance, guiding the choice of significant PCs for downstream clustering and dimensionality reduction

### 3.3. Ligand–Target Predictions

We next examined the predicted regulatory potential of ligands on target genes in the receiver population. The ligand–target heatmap (Figure 3) highlighted SIRPG as a central regulator, with predicted effects on genes such as CD47, CD5, and FCER1G. Other ligands, such as TNF and IFNG, were predicted to regulate genes associated with inflammatory signaling and effector functions, consistent with their known biological roles.

**Figure 3:**
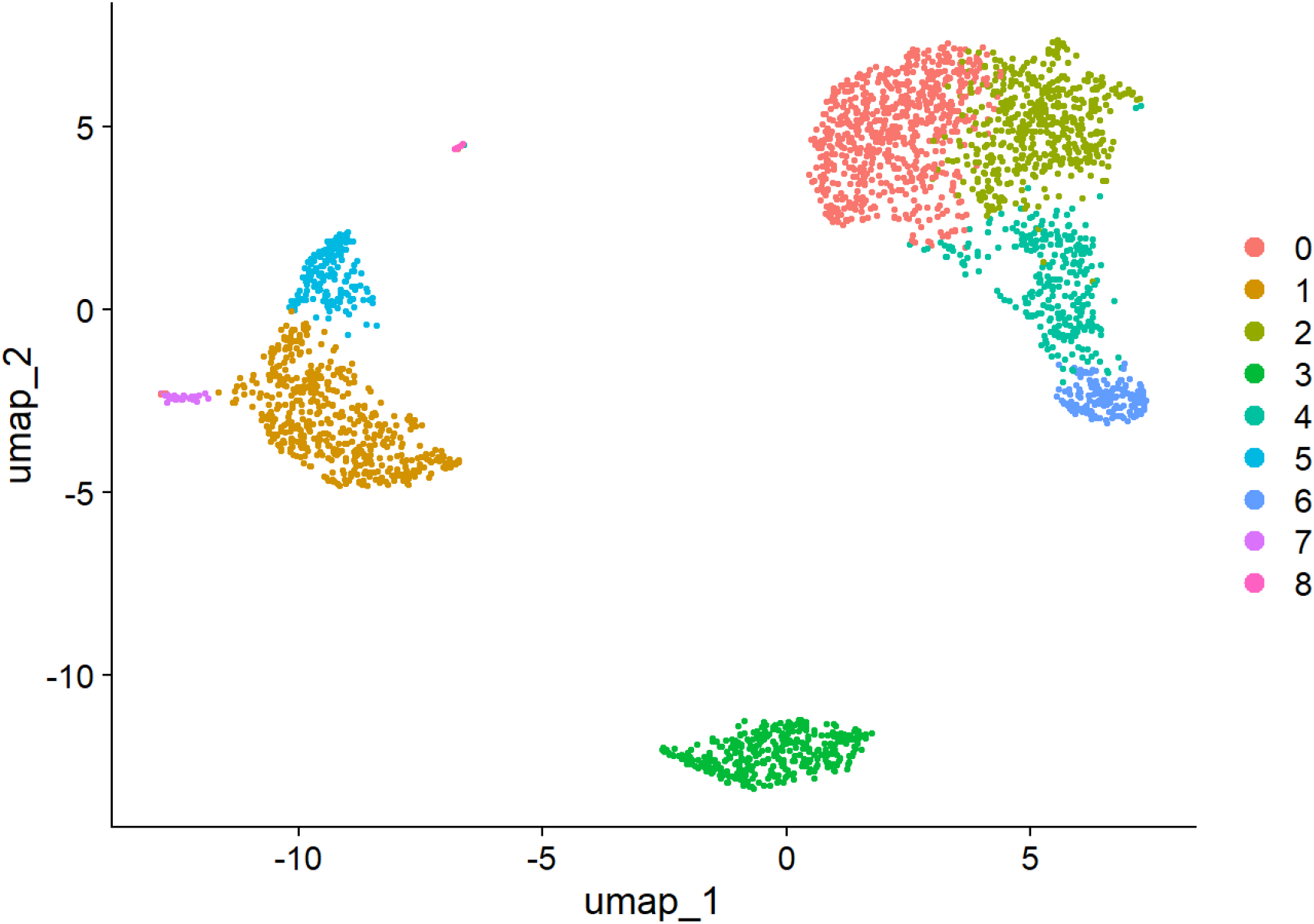
UMAP visualization of PBMC3k single-cell RNA-seq data after clustering. Each point represents a single cell, and cells are grouped into nine transcriptionally distinct clusters (0–8), highlighting the heterogeneity of immune cell populations in peripheral blood.

**Figure 4:**
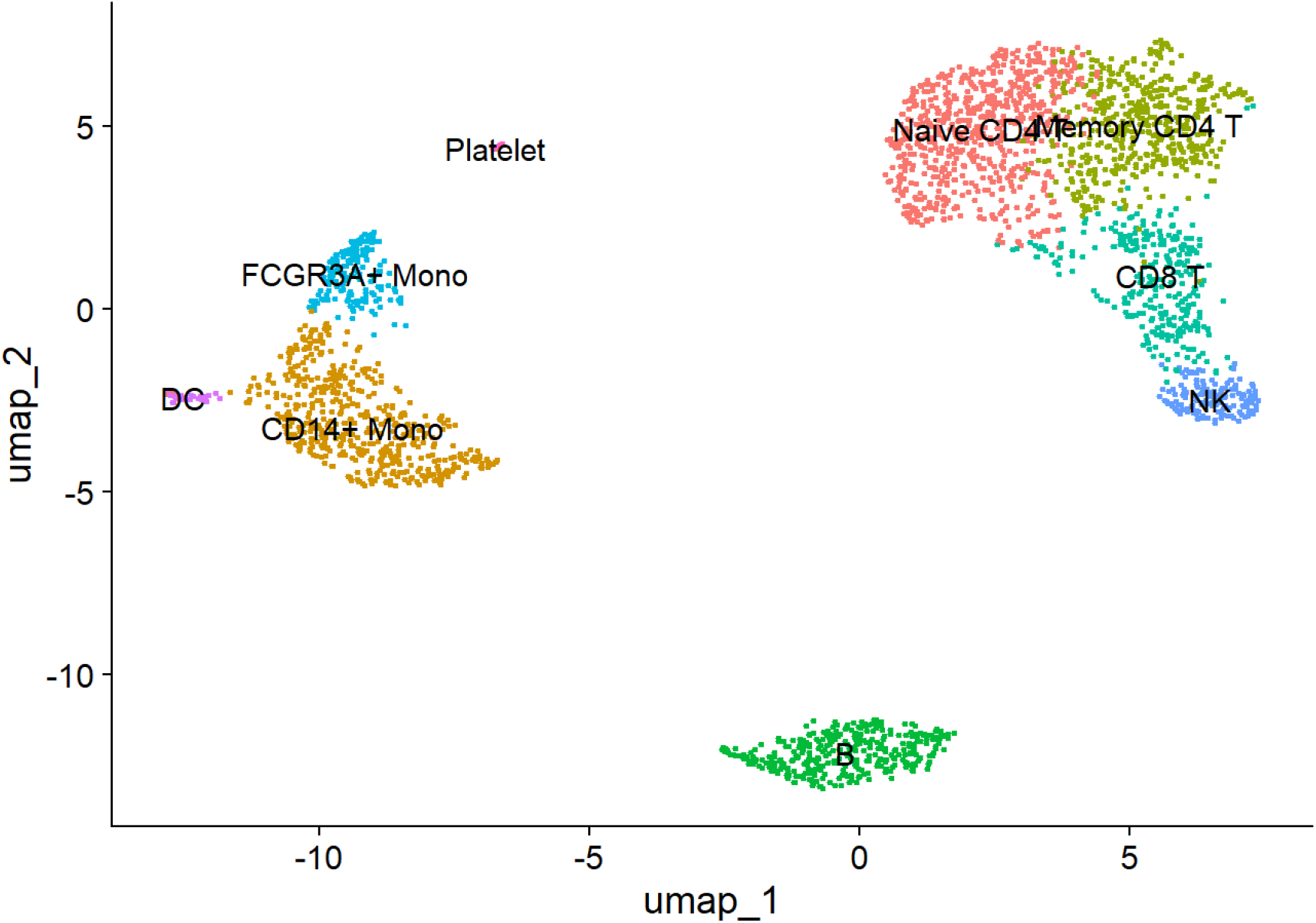
UMAP visualization of PBMC immune cell subsets. Uniform Manifold Approximation and Projection (UMAP) embedding of PBMC single-cell RNA-seq data after dimensionality reduction. Major immune populations were annotated based on canonical marker genes, including CD14^+^ monocytes, FCGR3A^+^ monocytes, dendritic cells (DCs), B cells, natural killer (NK) cells, CD8^+^ T cells, naïve CD4^+^ T cells, memory CD4^+^ T cells, and platelets.

**Figure 5:**
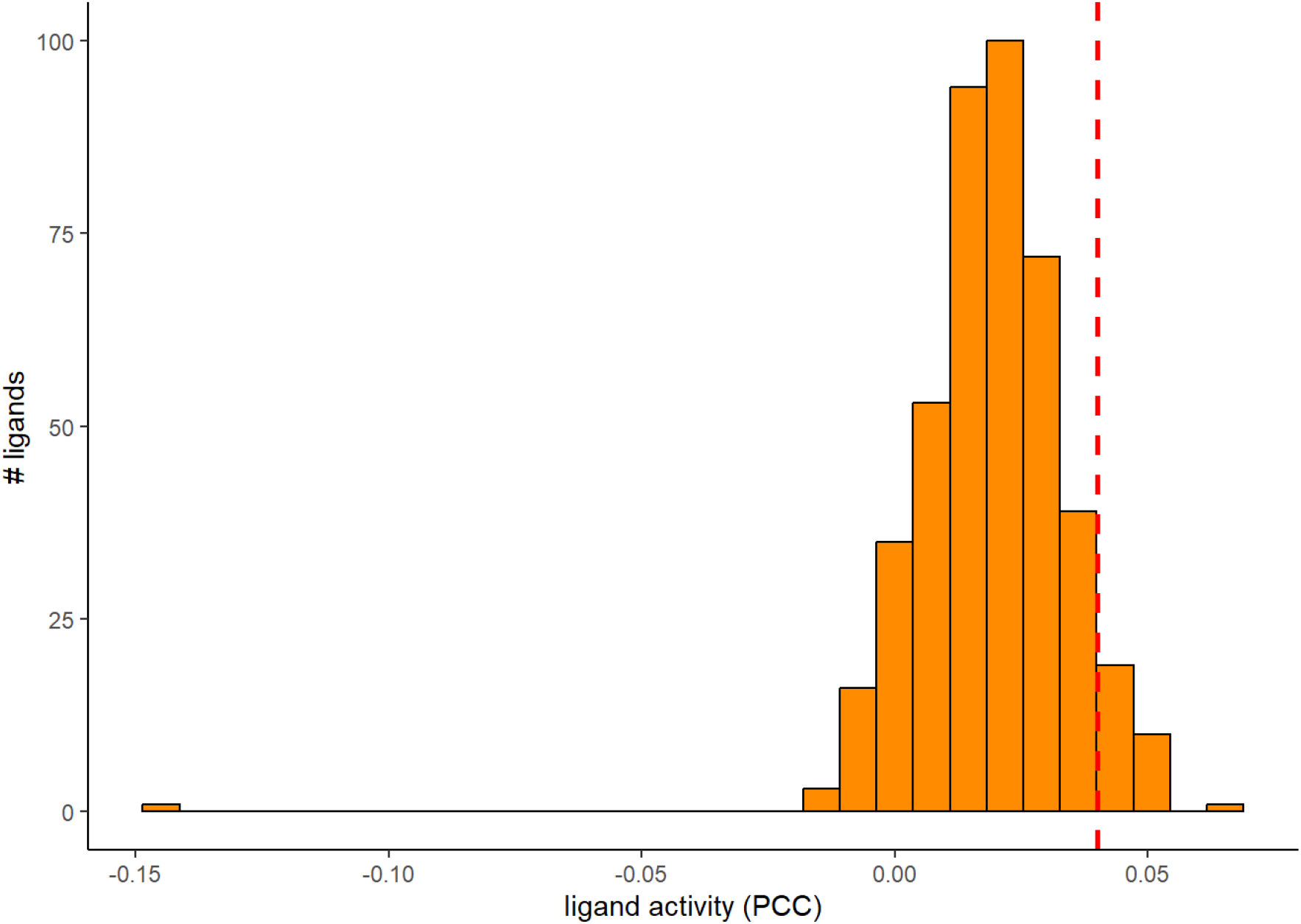
Histogram of ligand activity scores here

**Figure 6:**
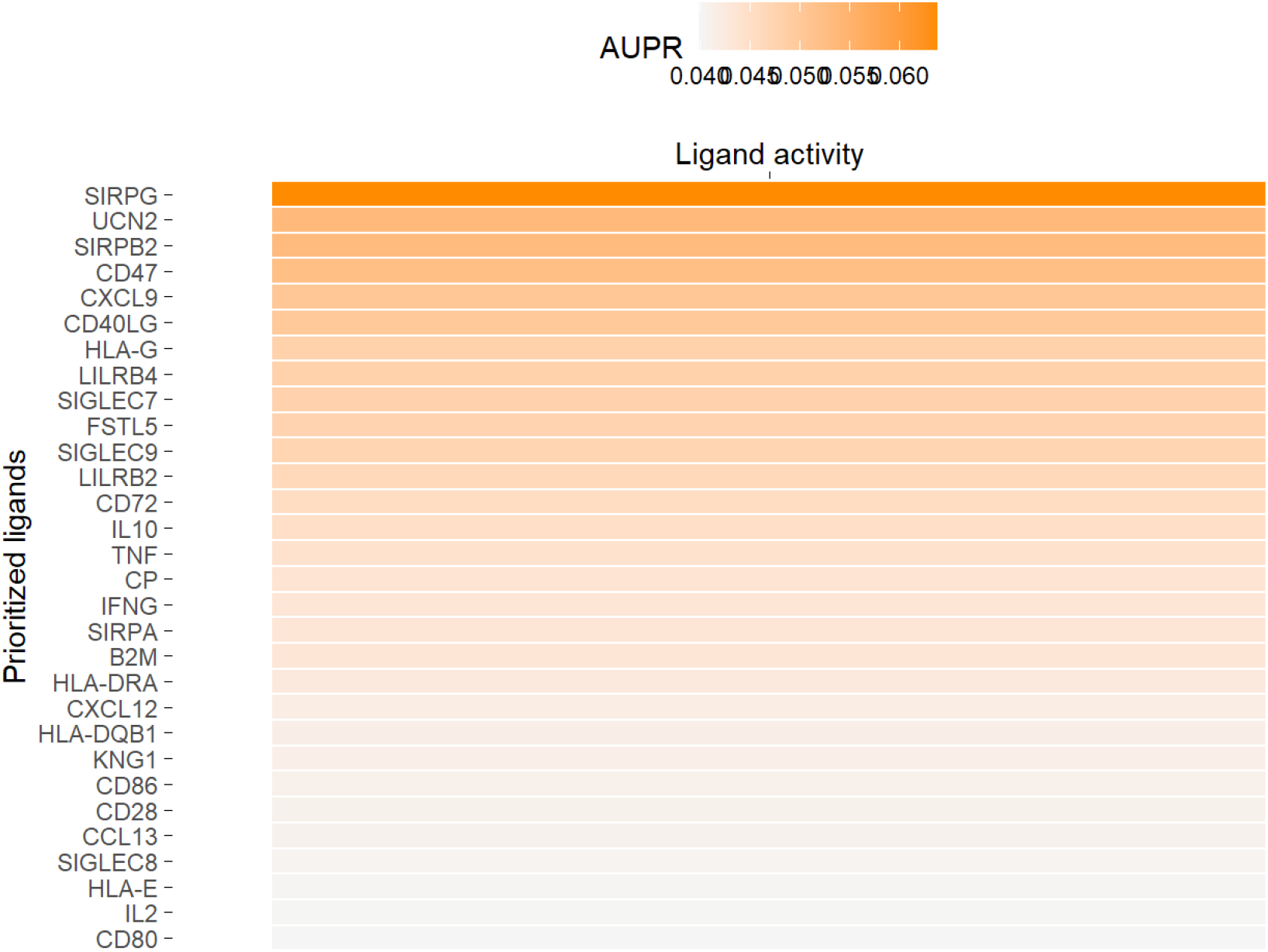
Heatmap of prioritized ligands and their ligand activity scores here.

**Figure 7:**
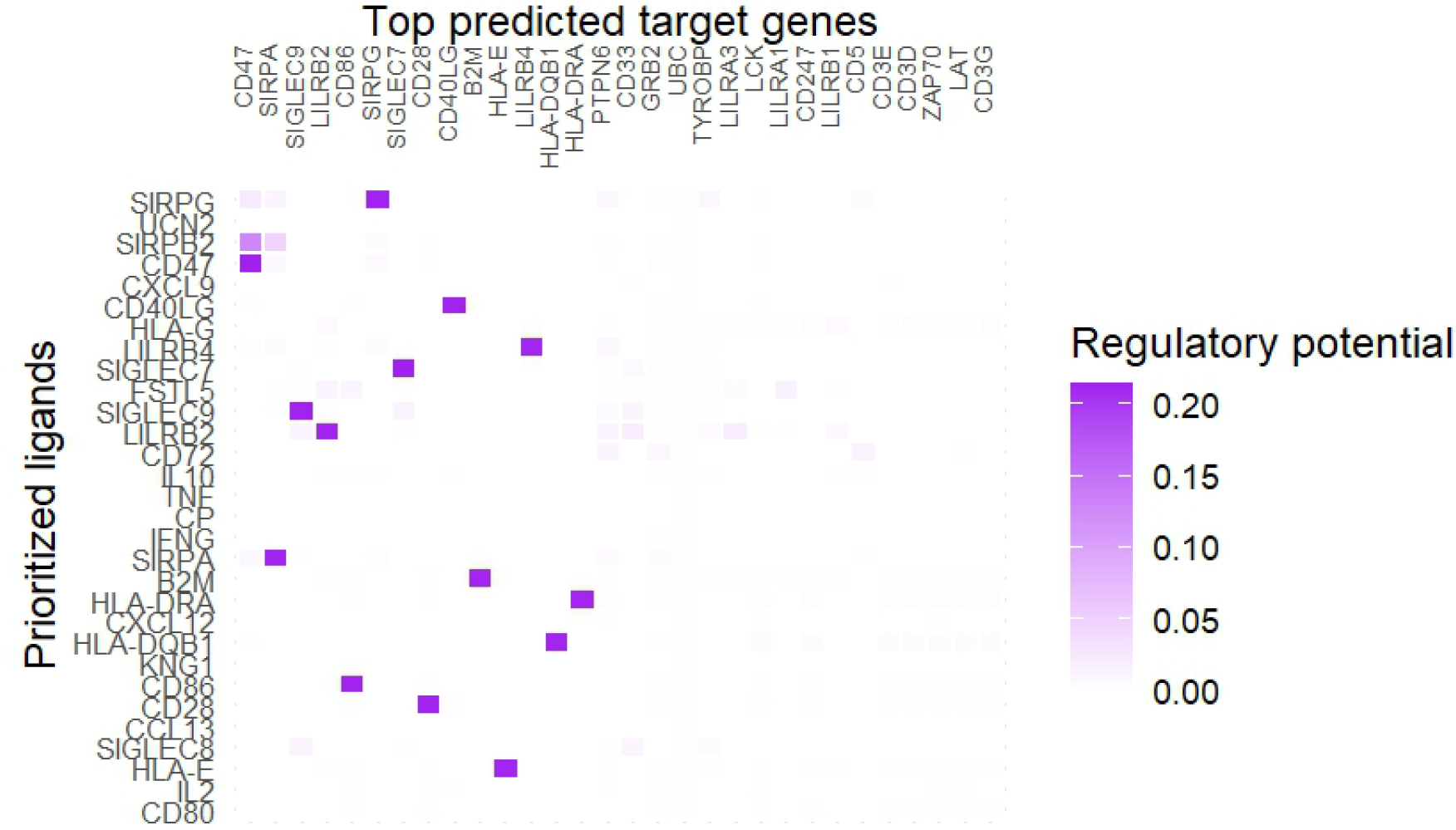
Ligand–target heatmap showing prioritized ligands versus predicted target genes.

### 3.4. Ligand–Receptor Interactions

Finally, mapping predicted ligands to expressed receptors on memory CD4^+^ T cells revealed key ligand–receptor pairs

**Table 1:** Predicted ligand–receptor interactions between CD14^+^ monocytes and memory CD4^+^ T cells.

| Ligand | Receptor(s) |
| --- | --- |
| SIRPG | CD47, CD5, CD9 |
| TNF | TNFRSF1A, TNFRSF1B |
| IL2 | IL2RB |
| CXCL9 | CXCR3 |

These findings highlight both classical cytokine signaling pathways and less-studied immunoregulatory axes (e.g., SIRPG–CD47), which may represent novel mechanisms of monocyte–T cell communication.

**Figure 8:**
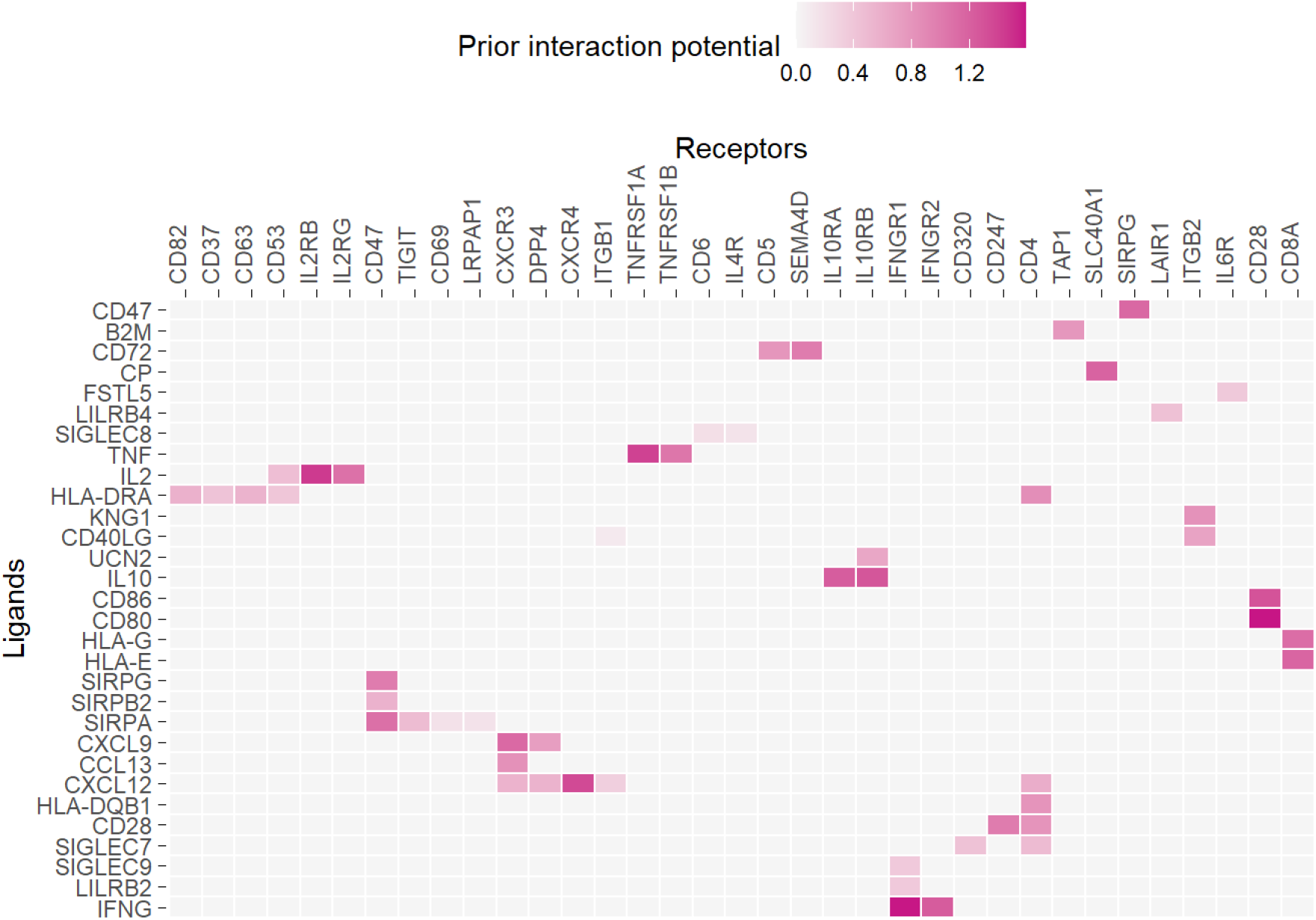
Heatmap of ligand–receptor interactions here

## 4. Discussion

Our NicheNet analysis of PBMC single-cell data revealed that CD14^+^ monocytes act as important modulators of memory CD4^+^ T cell responses through diverse ligand–receptor–target interactions. The results highlight both classical inflammatory pathways and less-studied regulatory axes that together contribute to shaping T cell transcriptional programs. Key predicted ligands included TNF, IL10, IFNG, and CXCL9, consistent with the well-documented roles of monocyte-derived cytokines in promoting T cell activation and effector function (Biron, 1999);(Viola and Lanzavecchia, 1996). The TNF–TNFRSF1A/TNFRSF1B axis, for example, is central to driving T cell activation and survival, aligning with our predictions (Watts, 2005). Similarly, IL2–IL2RB interactions are crucial for CD4^+^ T cell proliferation and differentiation (Boyman and Sprent, 2012). In addition to these classical pathways, our analysis identified the SIRPG–CD47 interaction as a top-ranked signal. SIRPG–CD47 interactions have been implicated in T cell adhesion and costimulation, and may represent an underappreciated mechanism by which monocytes modulate T cell responses (Yamaoka et al., 2020). This suggests that beyond cytokines, checkpoint-like interactions may also play an important role in shaping memory T cell function in peripheral blood. Taken together, these results support the concept that monocytes provide both inflammatory cues and regulatory signals to memory CD4^+^ T cells. Future validation of SIRPG–CD47 and related axes could yield novel insights into immune regulation, with potential therapeutic implications for diseases involving aberrant T cell activation such as autoimmunity and chronic inflammation.

## Code Availability

All code used for this analysis is available at: GitHub Repository: Pbmc Scrna seq Analysis

## Supplementary Figures

**Figure 9:**
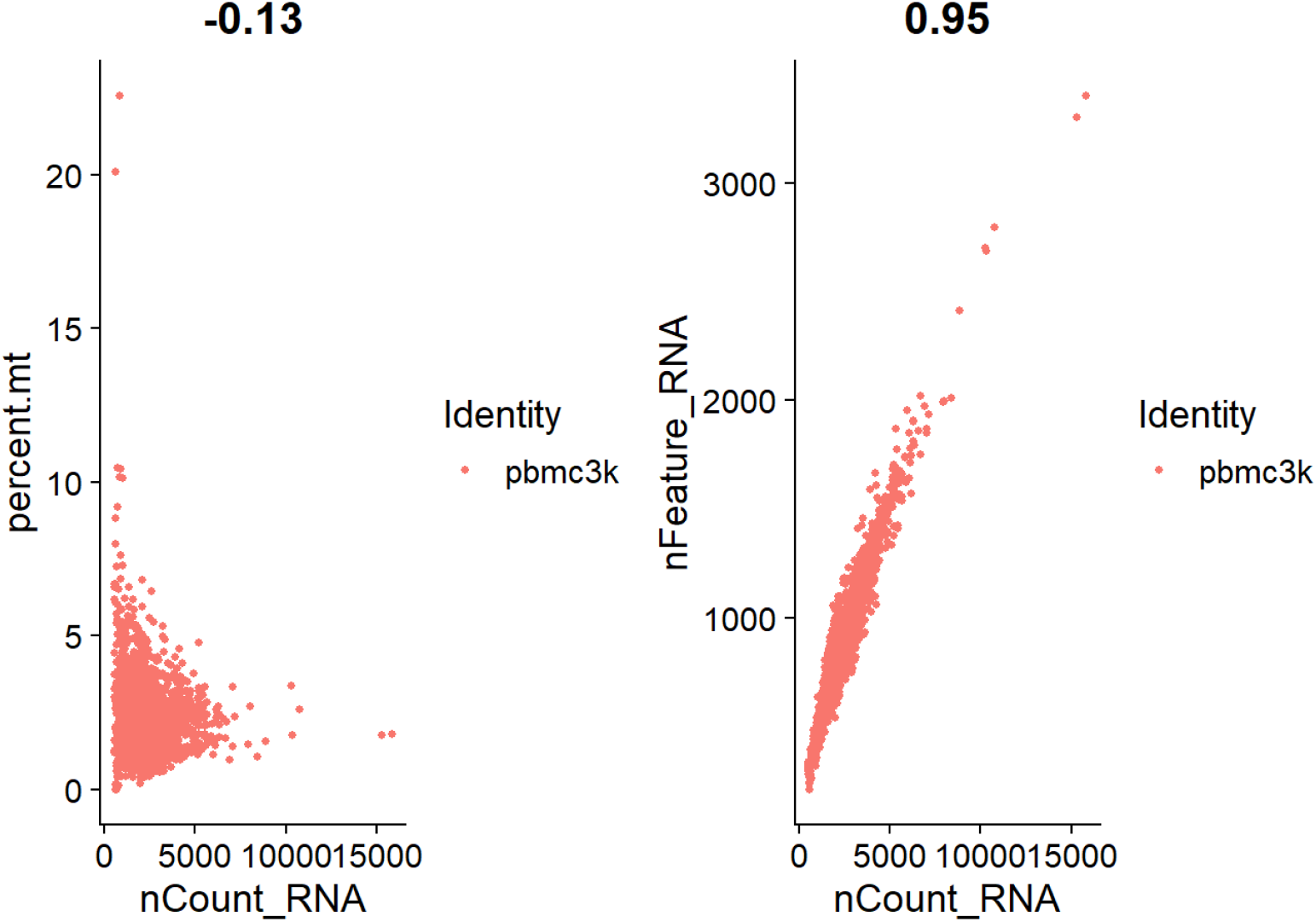
Scatterplots of quality control (QC) metrics for the PBMC3k dataset. (Left) Relationship between total RNA molecules per cell (nCount RNA) and the percentage of mitochondrial transcripts (percent.mt), showing a weak negative correlation (–0.13). (Right) Relationship between nCount RNA and the number of detected genes (nFeature RNA), showing a strong positive correlation (0.95). These QC metrics help identify low-quality or stressed cells for filtering.

**Figure 10:**
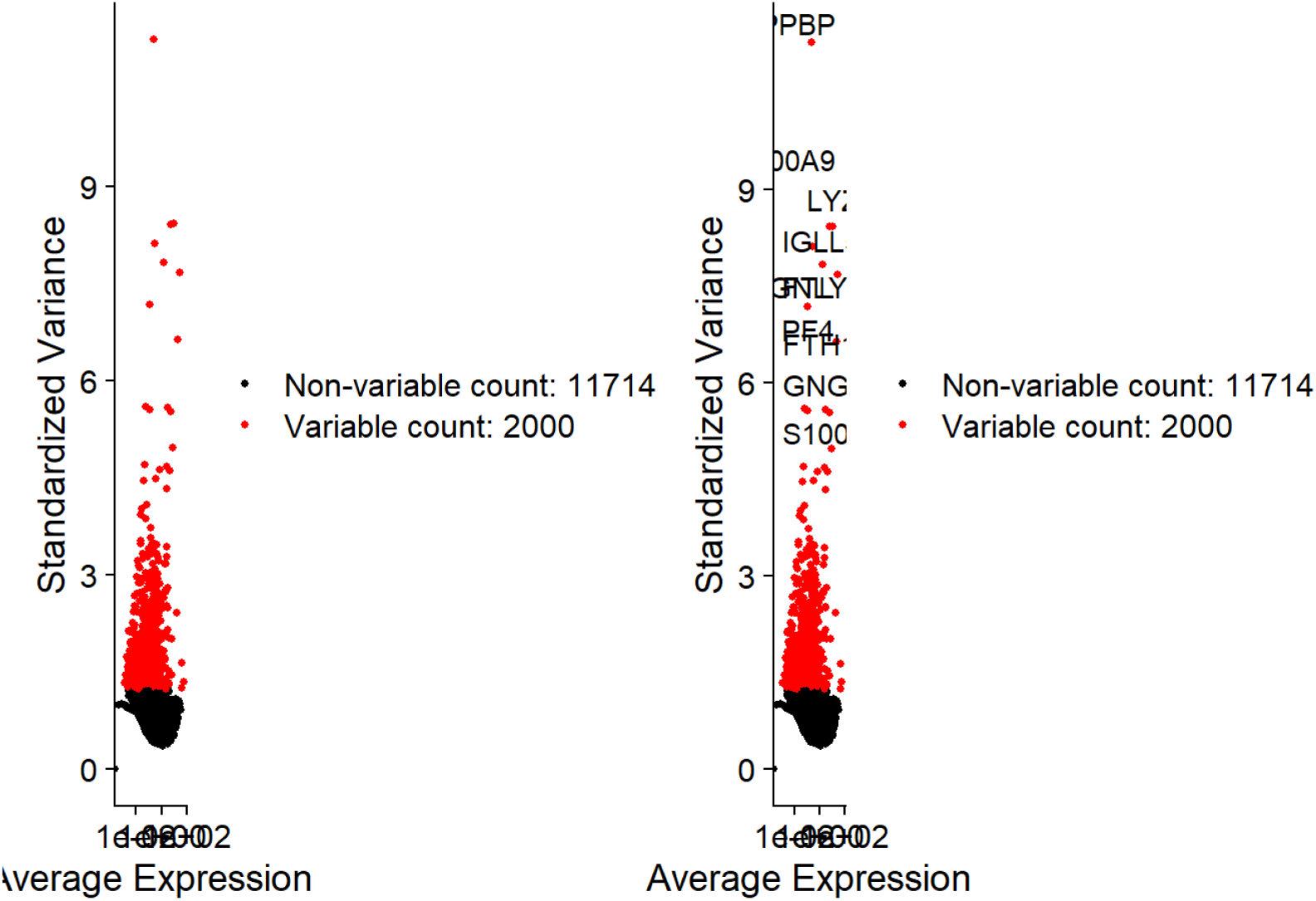
Identification of highly variable genes in the PBMC3k dataset. Scatterplots show standardized variance versus average expression. Black dots represent non-variable genes (11,714), while red dots indicate highly variable genes (2,000) selected for downstream analysis. The right panel highlights the top variable genes (e.g., *S100, LYZ, GNLY*) that contribute most to cellular heterogeneity.

**Figure 11:**
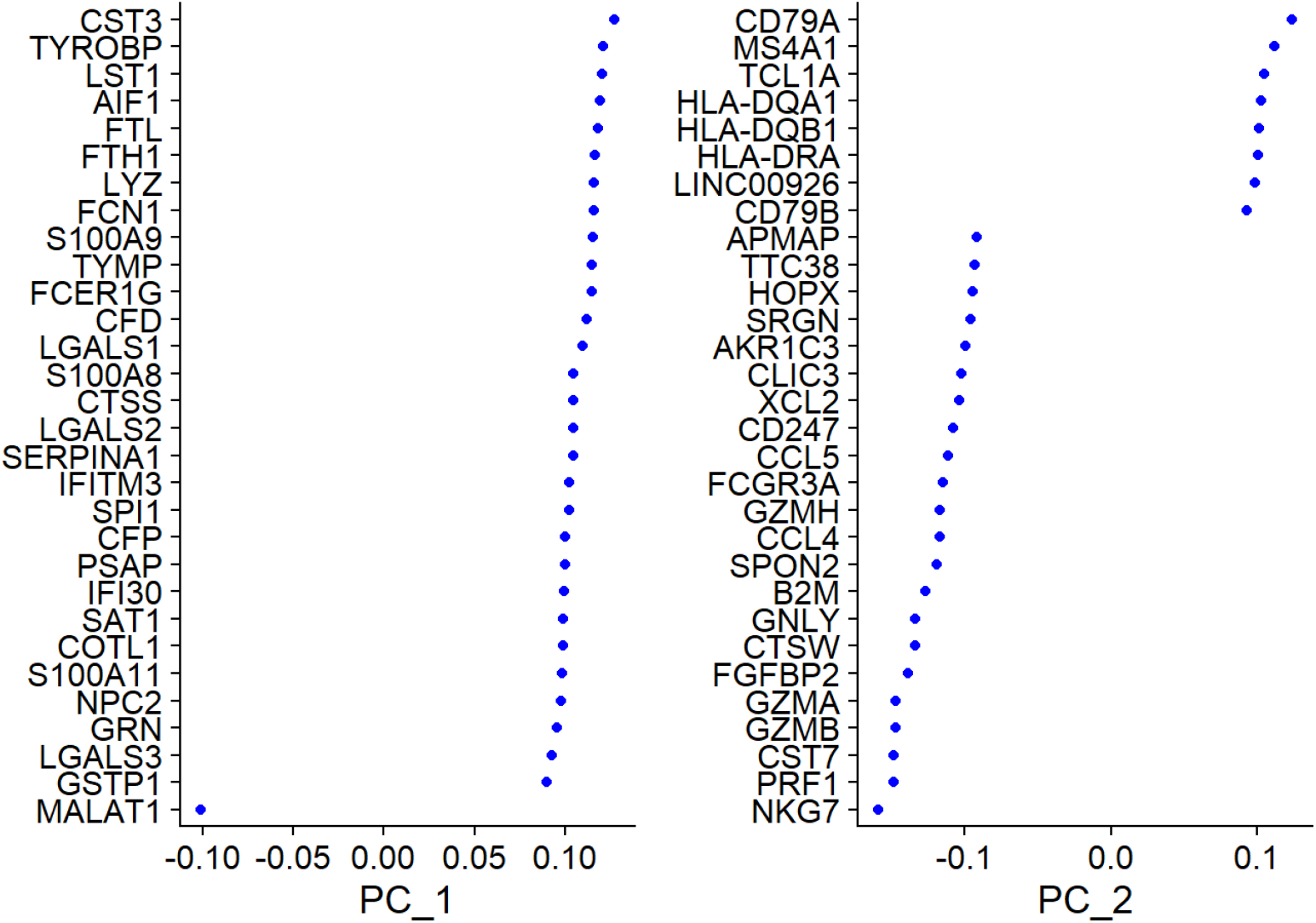
Principal component analysis (PCA) loadings plot showing the top genes contributing to PC1 and PC2 in the PBMC3k dataset. The left panel highlights genes with the strongest positive and negative loadings on PC1 (e.g., *CST3, TYROBP, LYZ*), while the right panel highlights genes contributing to PC2 (e.g., *CD79A, MS4A1, NKG7*). These genes drive the separation of major immune cell populations in the reduced dimensional space.

**Figure 12:**
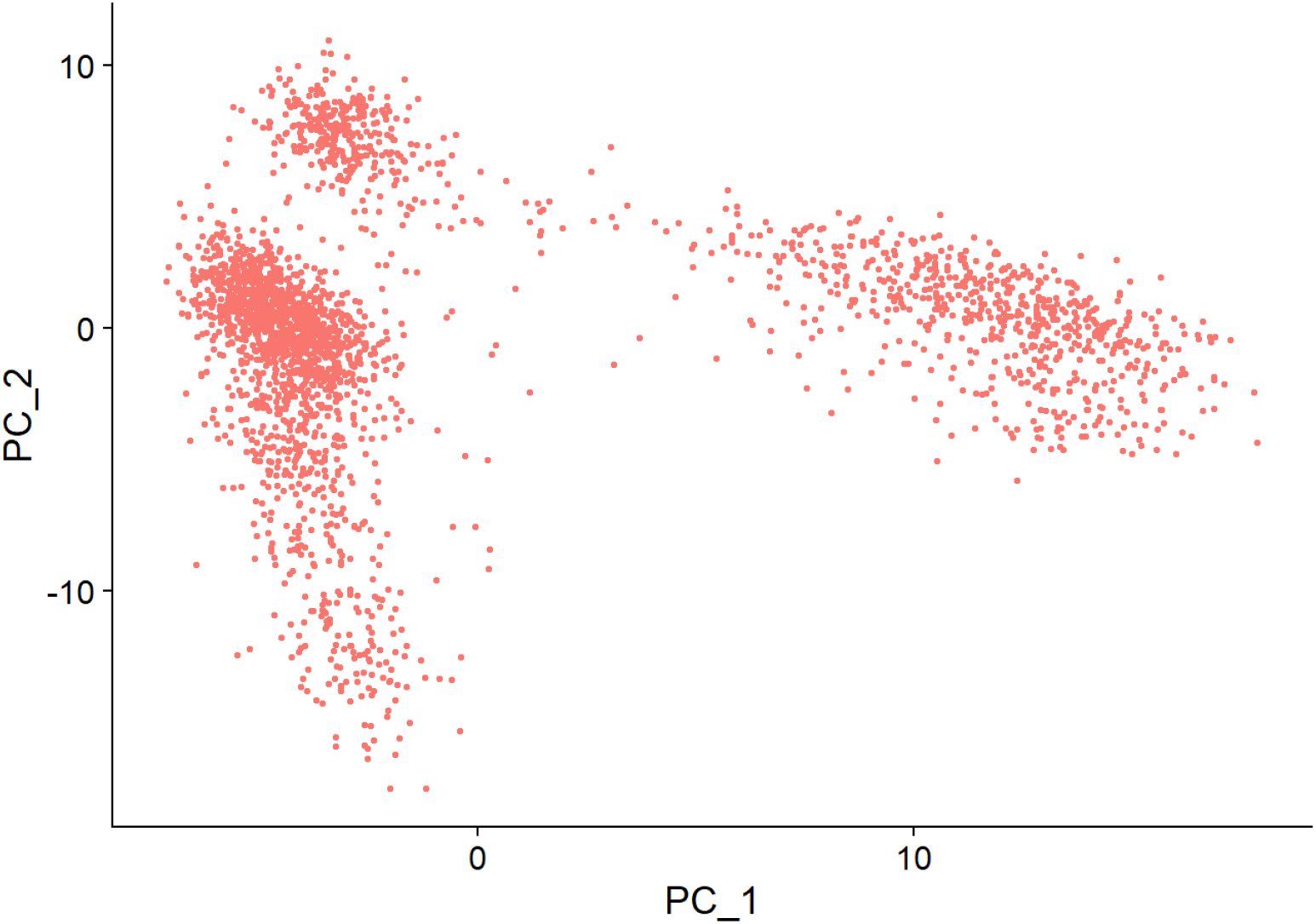
Principal component analysis (PCA) scatter plot of the PBMC3k dataset. Each dot represents a single cell projected into the first two principal components (PC1 and PC2). This visualization highlights variability in gene expression across cells and provides the basis for subsequent clustering and dimensionality reduction.

**Figure 13:**
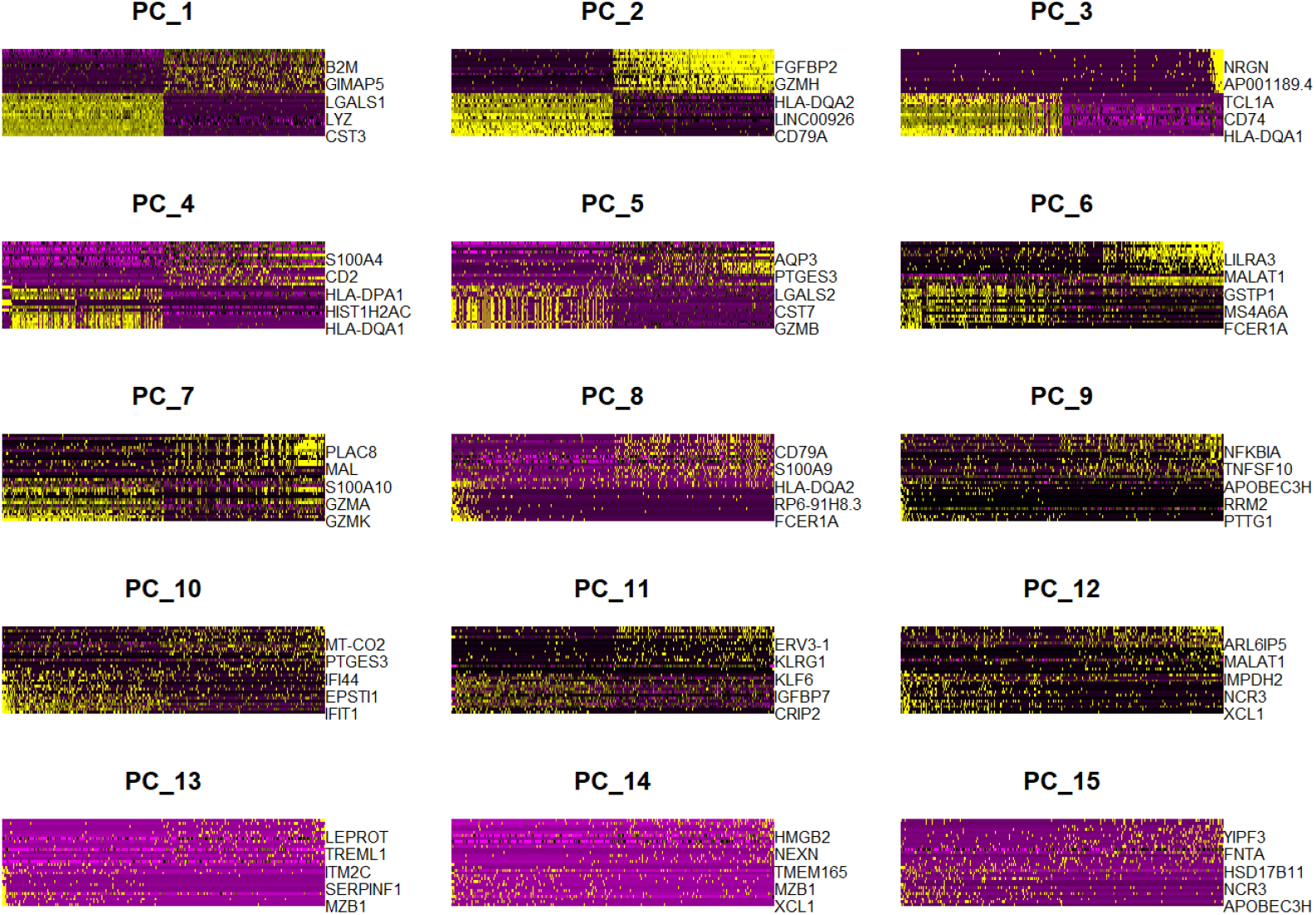
Heatmaps of the top principal components (PC1–PC15) from PCA on the PBMC3k dataset. Each panel shows the genes with the highest contributions to the corresponding principal component, with expression values color-coded (yellow = high, purple = low). These PCs capture major sources of transcriptional variability among cells and highlight key marker genes driving separation.

**Figure 14:**
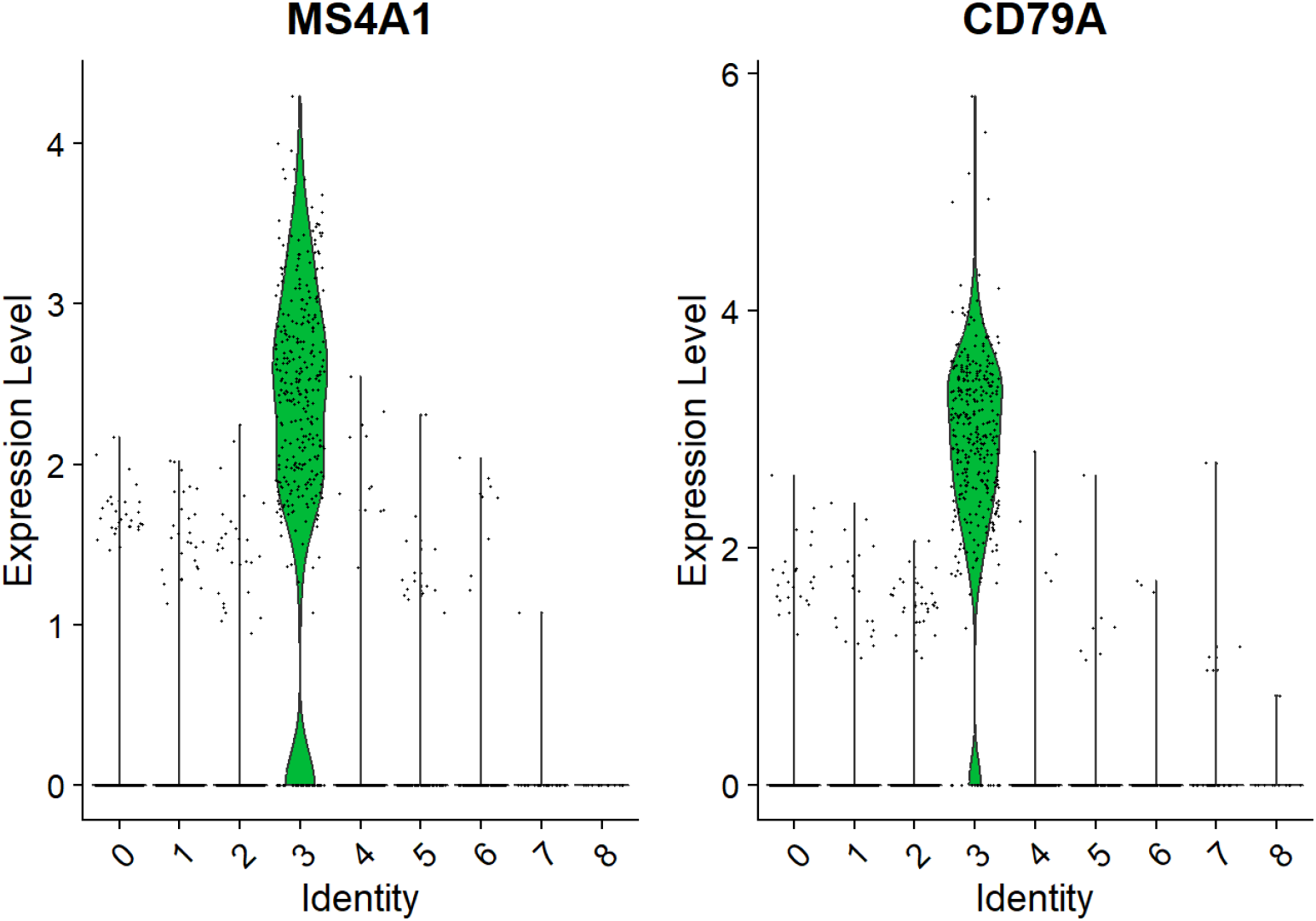
Violin plots showing expression levels of marker genes MS4A1 (left) and CD79A (right) across the identified clusters. Both genes exhibit strong expression in cluster 3, consistent with B cell identity, while showing minimal expression in other clusters.

**Figure 15:**
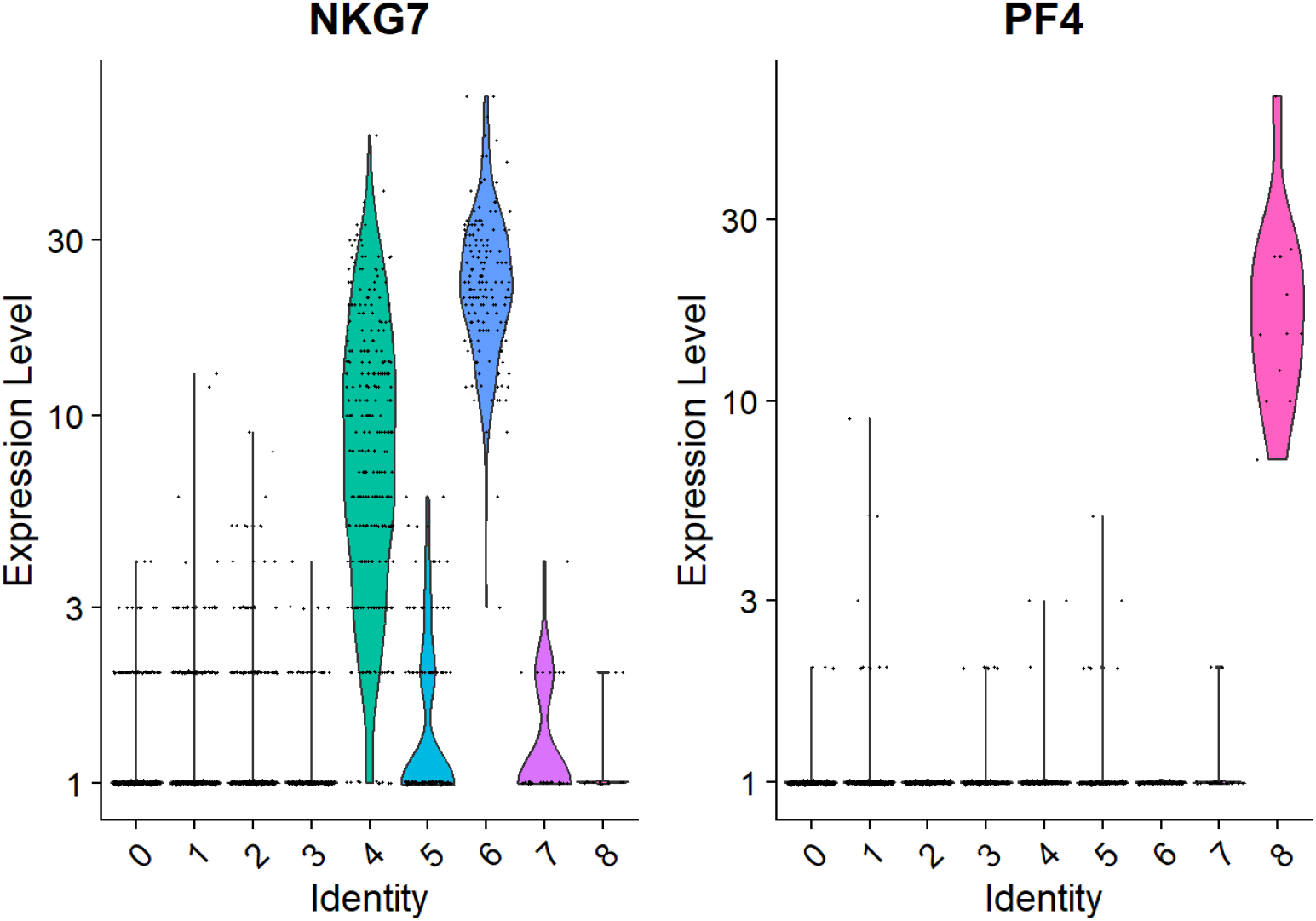
Violin plots showing expression levels of NK/platelet marker genes. **Left:** NKG7 expression is enriched in clusters 4 and 6, consistent with natural killer (NK) cells. **Right:** PF4 expression is highly enriched in cluster 7, consistent with platelet lineage identity.

**Figure 19:**
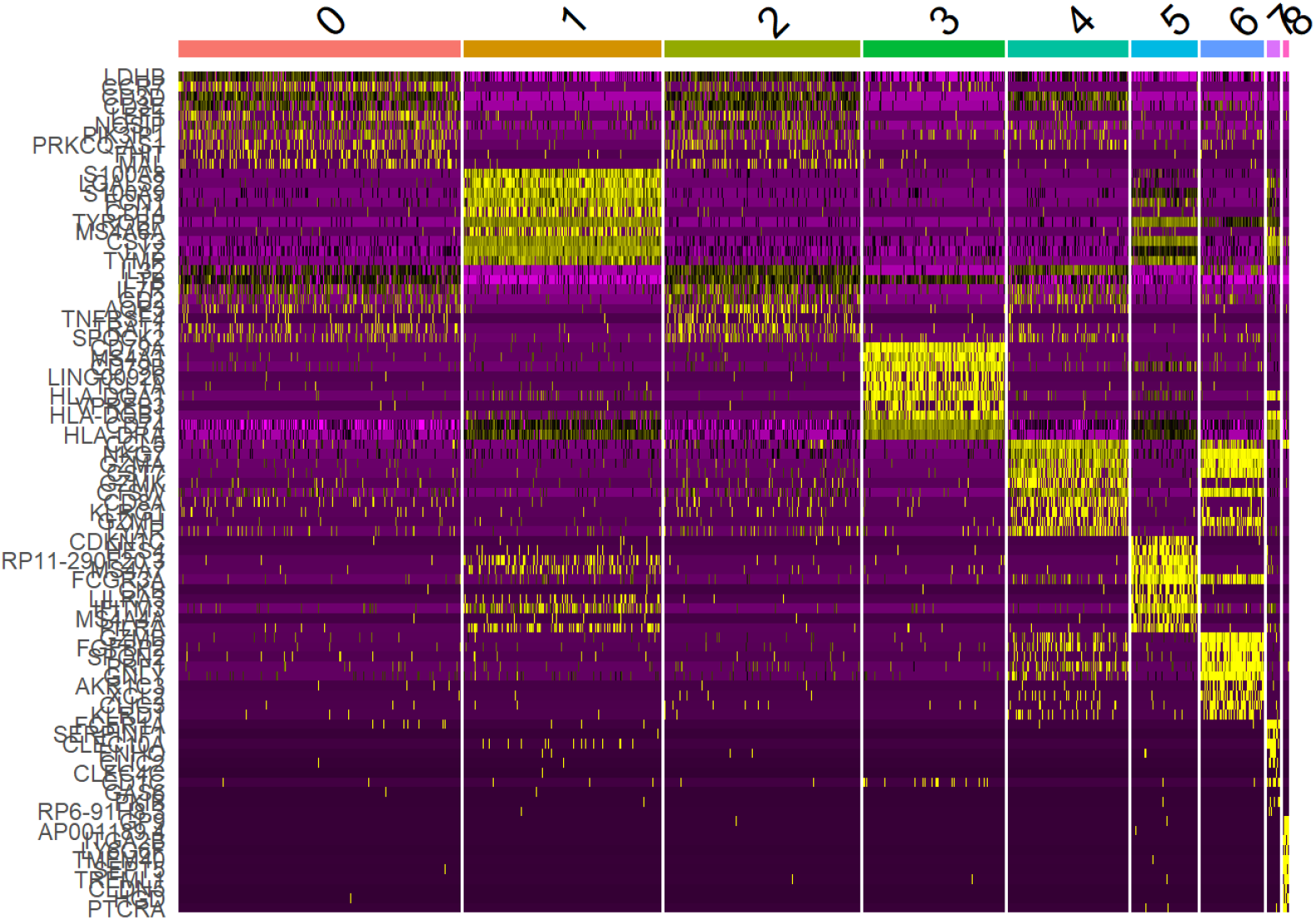
Heatmap of the top marker genes across clusters. Each column represents a cluster, and each row corresponds to a marker gene. Expression values are scaled and color-coded from low (purple) to high (yellow). Cluster-specific gene signatures highlight the distinct transcriptional identity of each cell type.

